# Exploring generality of experimental conformational changes with AlphaFold predictions

**DOI:** 10.1101/2022.04.12.488086

**Authors:** Albert Castellví, Ana Medina, Giovanna Petrillo, Theo Sagmeister, Tea Pavkov-Keller, Fernando Govantes, Kay Diederichs, Massimo D. Sammito, Isabel Usón

## Abstract

Structural predictions have matched the accuracy of experimental structures in the case of close homologues, outperformed docking methods for multimeric complexes and helped sampling the conformational landscape of transporters and receptors. Such successes prompt the question whether predictions can be used to relate experimental structures in the context of available knowledge. LysR-type transcriptional regulators (LTTR) constitute the most common family of bacterial regulators. Intriguingly, their experimental structures are remarkably diverse. The active species, composed of flexible monomers dimerizing through their N- and C-terminal domains in a circular arrangement, differ across LTTR, due to intrinsic sequence differences or because crystals stabilize diverse snapshots of a common dynamic mechanism. We have used AlphaFold2 (AF) to interrogate the experimental AtzR structure in the context of predictions guided towards the different hetero-multimeric conformations known for other LTTR. Our approach drives AF prediction with the structure-based selection of the information input through sequence alignment and template conformation, linked to examination of the energy with PISA and interactions with ALEPH.

## Introduction

LysR-type transcriptional regulators (LTTR) constitute the most abundant family of prokaryotic transcription regulators (Henikoff *et al*., 1988). Acting both as transcription activators and repressors, the mechanism underlying their activity involves specific interaction to chemical effectors, other proteins and their cognate DNA, requiring large dynamic changes in the active, typically tetrameric species (Maddocks & Oyston, 2008). Despite the difficulties to crystallise this kind of protein, several full-length structures have been determined, showing remarkable conformational diversity. In addition, numerous structures of LTTR C-terminal, effector-binding domains are available, reiterating a similar dimerization mode to that present in the full-length structures. The monomers are composed of a better conserved, N-terminal DNA-binding domain and a more variable C-terminal effector binding domain, linked by a hinge region mediating conformational variability. Two monomers in extended configurations and two in a bent, compact arrangement, with additional intramolecular contacts between both domains, typically coexist within a tetramer. Monomers associate into a ring, dimerising with different partners through their N-terminal as well as through their C-terminal domains. In many structures, additional contacts collapse the ring into a more compact particle. **Figure 1** displays representative cases of tetrameric conformations presenting pairs of exposed N-terminal, DNA-binding dimers. **Table 1** summarizes the 15 non-redundant tetrameric structures potentially apt for DNA binding, two structures where one or both DNA-binding dimers are buried and a rare octameric structure. **Table 1** is extended with the summary of structures containing the isolated effector binding domains. Only the full-length CbnR structure has been determined in complex to a region of its DNA promoter (PDB ID 7D98, Giannopoulou *et al*., 2021) and generally, ambiguity about activation state lingers even in the presence of effector, given the rich stabilisation buffer required to avoid aggregation (Monferrer *et al*., 2008) and crystal constraints. For instance, no conformational differences are seen in TsaR upon ligand binding (Monferrer *et al*., 2010) and the CbnR DNA-bound structure evidences no overall change versus the unbound protein (PDB ID 1IXC Muraoka *et al*., 2003). Therefore, the question of relating all the available structural information on LTTR to a common frame remains unsettled.

**Table 1.**
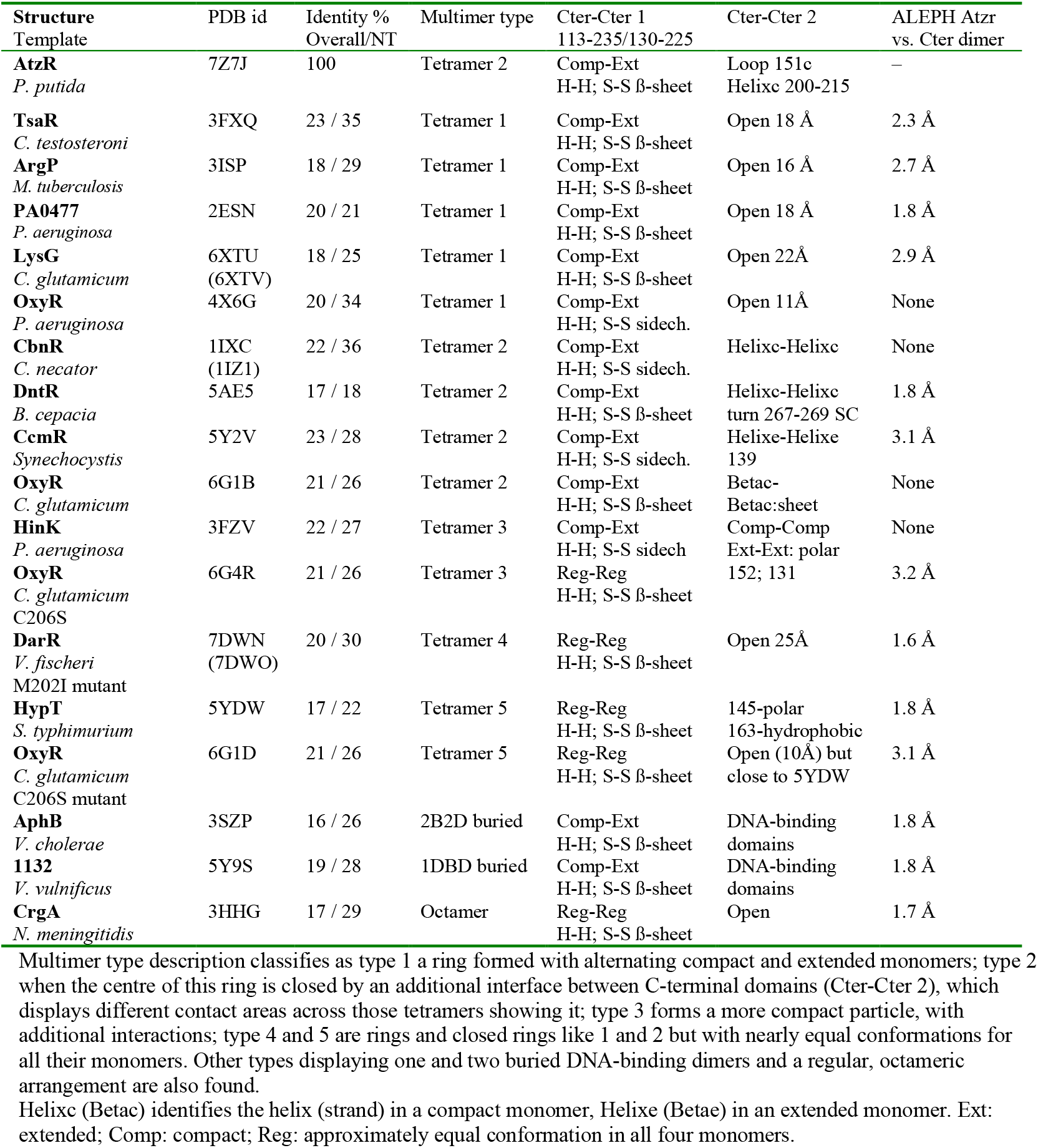
Summary of LTTR full-length crystallographic structures used as templates in predictions, ordered by tetrameric conformation as in Figure 1. Analysis of main interfaces and contacts undergone by the effector binding domains.

**Figure 1.**
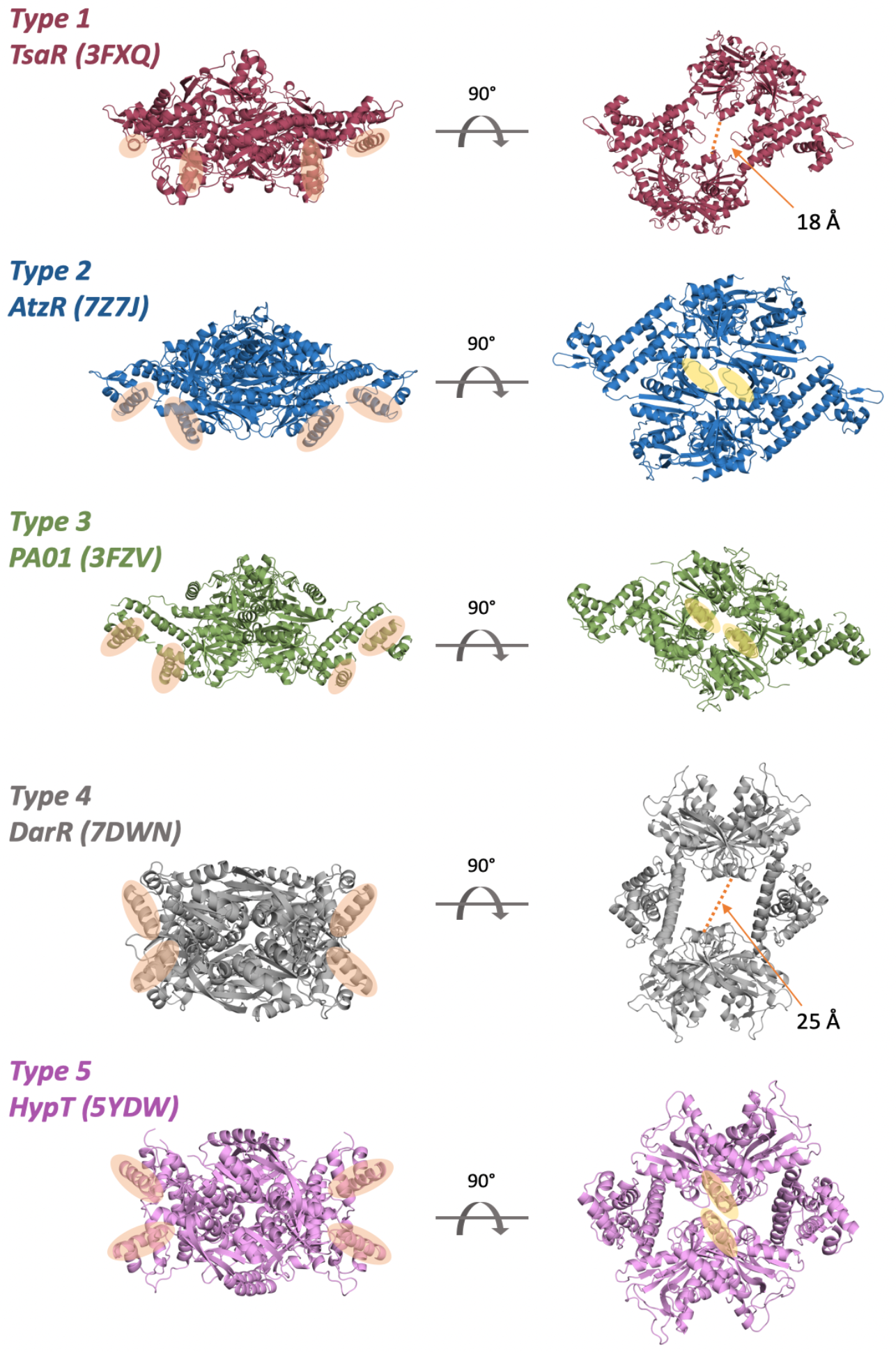
Representative conformations displayed by full-length LTTR. Side and top view for different experimental structures categorised as tetramers type 1 to 5. Orange circles highlight the DNA-binding helices (left). Yellow circles highlight the interacting areas that close the central ring (right).

AlphaFold (AF) (Jumper *et al*., 2021) has brought the accuracy of sequence-based structure predictions to a level of atomic detail comparable to that derived from close homologs, as seen in CASP13-14 (Kryshtafovych *et al*., 2019). This quality is verified through the stringent test of overcoming the crystallographic phase problem as the new AF and RoseTTaFold models are accomplishing the solution of crystallographic structures through molecular replacement (Baek *et al*., 2021). AF application has been extended to the prediction of multimers (Evans *et al*., 2022), reaching a far superior success in the case of heterodimers than state of the art docking or fold and dock methods (Bryant *et al*., 2021). The information exploited in the structural prediction for a given sequence relies on the alignment to homologs and the pairwise conservation of residues involved in contacts (Marks *et al*., 2011). It has been shown that modifying the shallowness of the input sequence alignments can be used to sample the conformational landscape of transporters and receptors in monomeric structures adopting multiple states in a dynamic mechanism (del Alamo *et al*., 2022). The power of AF to bring into the predictions broad prior knowledge learned from all known structures and sequences unrelated to the target and flexibly evolve beyond the nearest information suggests a potential use emulating classical homology-modelling (Sali & Blundell, 1993; Waterhouse *et al*., 2018; Zheng *et al*., 2021). In the case of the LTTR structures, two different conformations for the monomers –extended and compact– typically coexist in the LTTR multimers and undergo different interactions. Interconversion of structural types may take place. Thus, individual chains do not simultaneously satisfy all restraints informed by conservation and even though they share a common sequence, structurally most of them resemble heteromultimers. This prompted us to assess the prediction of a new LTTR in AF, tailoring the input in order to drive the answer towards the particular association seen in each previously reported LTTR and to query the available experimental information in partial structures through specific local folds with ALEPH (Medina *et al*., 2020).

In the present work we have determined the new crystallographic structure of AtzR (Porrua *et al*., 2007) and interpreted it in the context of the previous experimental knowledge on LTTR systems, using AF to formulate the following questions: Can AtzR conformations be predicted to match all other previously described LTTR multimer? Can potential functionally relevant structural states of AtzR be predicted by AF, based on previously described LTTR oligomers? Can the feasibility of the predicted multimers be rated through the stability of the resulting interfaces? Will classification of the results inform the question whether dynamics are generally shared within the LTTR family or unique to the different members or within sets of them? To address these questions, the structural landscape of experiments and guided predictions has been interpreted using ALEPH (Medina *et al*., 2020) to characterise and compare interactions within the multimers and PISA (Krissinel, 2011) to estimate interface energy.

## Results

The structure of AtzR was crystallised in the presence of its effector, cyanuric acid (Porrua *et al*., 2007) and determined with ARCIMBOLDO_SHREDDER (Millán *et al*., 2018) using AF models. The asymmetric unit contains two monomers: one in extended and one in a compact conformation (**Figure 2a**), generating by symmetry the tetramer displayed in **Figures 1** and **2b**. As in other LTTR, monomers are composed of an N-terminal domain encompassing the first 90 amino acids building a winged helix-turn-helix motif in its first 60 residues and a nearly thirty amino acids long linker helix. The remaining 210 amino acids build the effector binding domain containing two Rossman-like subdomains with the binding site between them. The long linker helices in the N-terminal domains associate pairwise in an antiparallel disposition, presenting the DNA-binding motifs at a distance suitable for interaction with two consecutive major grooves. Both DNA-binding dimers are exposed on the same convex surface of the tetramer. Also the C-terminal domains dimerise through an interface built mainly by the association of two pairs of helices and the association of two pairs of beta strands through their main chain, resulting in the extension of a beta sheet (**Figure 2, Table 1**). The tetrameric ring is tightly closed by an extended contact interface between the C-terminal domains of two monomers in compact conformation. This contact interface between both domains is formed by 24 residues comprising residues 151-157 and the helices 197-214 in both chains. The electron density map showed clear density (**Figure 2c**) for cyanurate, the effector known to activate AtzR-mediated transcription. It binds to four regions, being coordinated by residues Ala101, Asp129, Ser200, Gly201, Gln241 and two water molecules (2 and 20) and stacked between Tyr225 and Phe148. Ser200 and Gly201 are located at one end of the helix involved in the ring-closing interface and Phe148 next to the loop at the centre of this interface.

**Figure 2.**
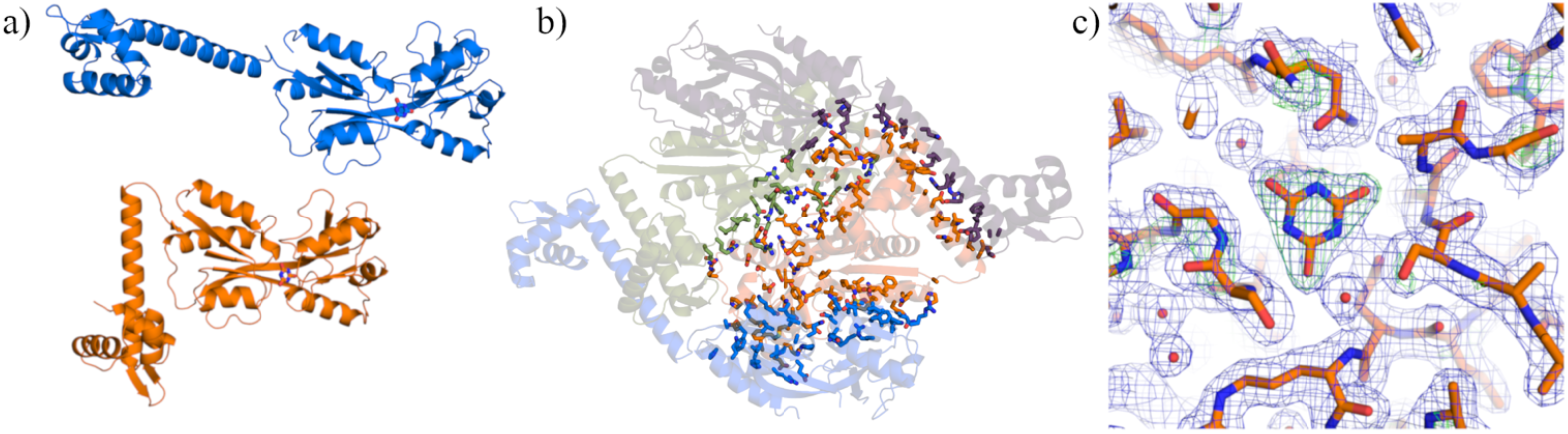
Experimental structure of AtzR. **a**) AtzR, as many other LTTR is composed of two monomers in an extended conformation and two in a compact conformation, with the hinge between the DNA-binding and the effector binding domain. **b**) The three different interfaces in the AtzR tetramer are highlighted showing as sticks the side chains of the residues involved. **c**) Omit at 2.5σ (green) and sigmaA-weighted 2Fo-Fc at 1σ (blue) electron density maps in the area of the cyanurate effector and its environment for chain A.

The other known tetrameric full-length structures were analysed, evaluating the interfaces built between monomers. The overall conformation of the tetramer is determined by the linker mediated angles between monomer domains, entailing corresponding differences in the interfaces. Attending to these interfaces, a classification of the tetrameric structures is summarised in **Table 1**. All tetrameric structures form rings built by dimerising N-terminal domains through the association of their long linker helices in antiparallel fashion and between pairs of C-terminal domains through symmetric pairs of helices and beta strands. This last interaction is either an extension of the beta sheets in the Rossman fold subdomains, formed through mainchain hydrogen bonding -as in AtzR- or an interaction through the side chains of these same strands. Intermediate situations, with paired strands in one side of the interface and side-chain bound ones in the other also arise, e.g. in CmpR. In contrast, the dimerisation of N-terminal domains is rather constant, leading to the presentation of two DNA-binding winged helix-turn-helix motifs, spaced to the distance between two major grooves as seen in the structure of the N-terminal domain of CbnR bound to DNA (PDB ID 5XXP, Koentjoro *et al*., 2018). This set of interactions builds tetramers classified for the purpose of this study as type 1 (**Table 1, Figure 1**). Additional interactions between C-terminal domains may close the ring (type 2), as in AtzR or CbnR, or build more compact particles as in type 3 seen in HinK (PDB ID 3FZV, Wang *et al*., 2021). Concomitantly, the distance and relative orientation between both pairs of DNA-binding dimers is different in all types of tetramers. More regular open and closed rings, with (nearly) equivalent conformations in their four monomers are seen in types 4 (DarR) and 5 (HypT). To aid our classification and calibrate our method (**Figure 3**), the experimental AtzR dimerization interface for the C-terminal domains was compared with that found in the other full-length structures in the PDB. Evidence for the interfaces formed in the tetramers was also queried against the experimental structures of the effector-binding domains with ALEPH (**Table 1 Extended, Figure 3**). The template used to represent the interface of AtzR comprised the central residues of each of the secondary structure elements involved in the interface, four helices and four strands. A structurally close arrangement matching this template was found in all full-length structures except for the ones where the main chain interaction between pairs of beta strands is substituted by a more distant interaction involving side chains and in an oxidised form of OxyR (PDB ID 6G1B, Pedre *et al*., 2018), involving the formation of a disulfide bridge causing a rearrangement of secondary structure, notably the alpha helix present in the reduced form is substituted by a pair of beta strands. The interface queried was also identified wherever present, in many of the effector-binding domain dimers. Some structures do not contain such dimers and others are distorted by a relative rotation between both pairs of Rossman fold subdomains.

**Figure 3.**
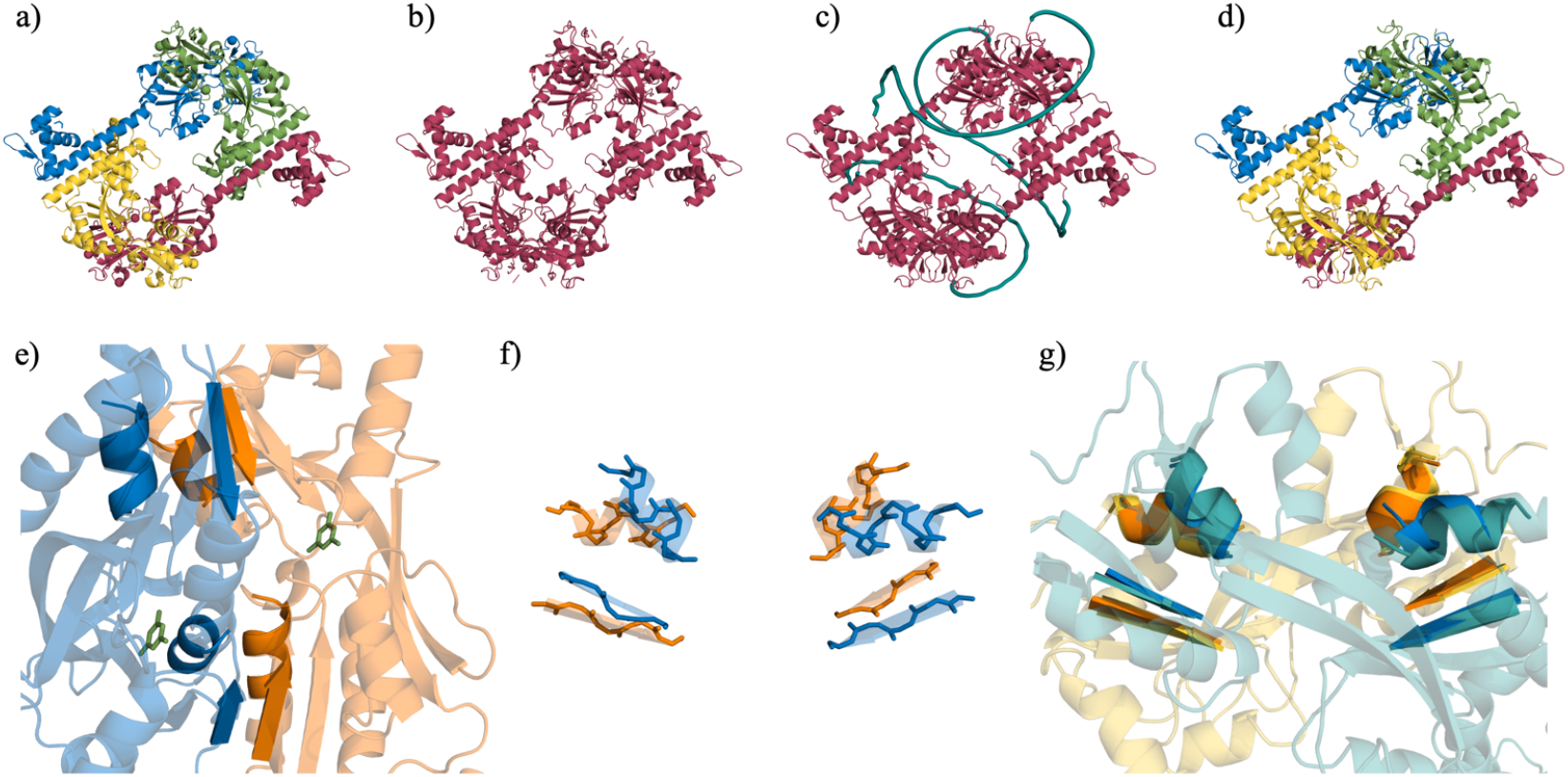
Methods for LTTR tetramer prediction and analysis. **a**) A single experimental tetramer is taken as template, and in the alignment to the target sequence. **b**) poly-glycine linkers are inserted in the sequence and the template structure is renumbered as a single chain. **c**) Resulting AF prediction with the unstructured glycine linkers (teal) **d**) The model is split in chains, renumbered and linkers are eliminated. Each chain is displayed in a different colour. **e**) The AtzR secondary structure elements involved in the dimer interface between effector-binding domains. **f**) Central amino acids for 4 helices and 4 strands are used to characterise the interface. **g**) Identification of the equivalent interface in other LTTR structures is performed in ALEPH.

To recreate possible structures of AtzR in the other conformations seen in the case of its homologs, in case that all or some of them might represent snapshots in a common dynamic mechanism, we used AF to predict structures targeting the different conformations. Our approach, implemented in ARCIMBOLDO_AIR, guides the AF prediction bypassing its native search and substituting it by designed input features. Here, we are providing structurally homogeneous information through alignment and template to calculate structures for a given sequence resembling the different multimeric conformations. In the present case, the sequence was that of AtzR, whose structure was absent from the databases and we strived to obtain its structural versions for each of the 16 tetramers in **Table 1**.

Our first experiment was to predict structural analogues for the different monomeric conformations seen in all these tetramers, assemble the predicted monomers most closely resembling those seen in the experimental structure (as judged by the rmsd of all Cα) into the corresponding tetramer and use the resulting template for a new prediction of the multimer as a monomeric chain. In eight of the cases, notably the type 1 and 2 tetramers, close conformations were obtained, within 6Å rmsd for a Cα superposition of template monomer and prediction. We observed that deviations from ideal geometry in the templates led to elongation of the regular secondary structure in the prediction. For the rather unconstrained, extended monomers deviations in the linker angle were large and influenced by the idealisation of the linker residues. Even in the best cases, stereochemistry of the resulting tetramer would be too poor for energy calculations to be useful and no further adjustment would take place in a subsequent prediction round when using a template of identical sequence to the target. The resulting models showed clashes and unlikely stereochemistry, rendering energetical or structural assessment unfeasible at an atomic level and were abandoned.

Instead, as illustrated in **Figure 3**, multimers annotated as a single chain with gaps corresponding to linker regions were generated from the 16 LTTR experimental structures and used as templates, limiting the sequence alignment to that of the template and four copies of the AtzR sequence with poly-glycine stretches inserted between sequence copies (Evans *et al*., 2022). Poly-glycine stretches are unstructured in the prediction but avoid distortions in the monomers they link. All conformations corresponding to the 16 tetramers were obtained for the AtzR sequence within rmsd values ranging from 2.7 to 5.3 Å for core Cα atoms of the tetramer versus template (**Table 2**). An increase in the percentage of ideal alpha-helical or beta-strand secondary structure in the predicted models versus their templates by 2-7% is noticeable. The secondary structure content in the predictions is always higher than the 68% characterising the experimental AtzR structure (**Table 2**). The stereochemistry for these predictions was favourable enough to assess them through the estimated binding energy (ΔGint) and the free Gibbs energy of dissociation of their interfaces as well as atomic structural tools.

**Table 2.**
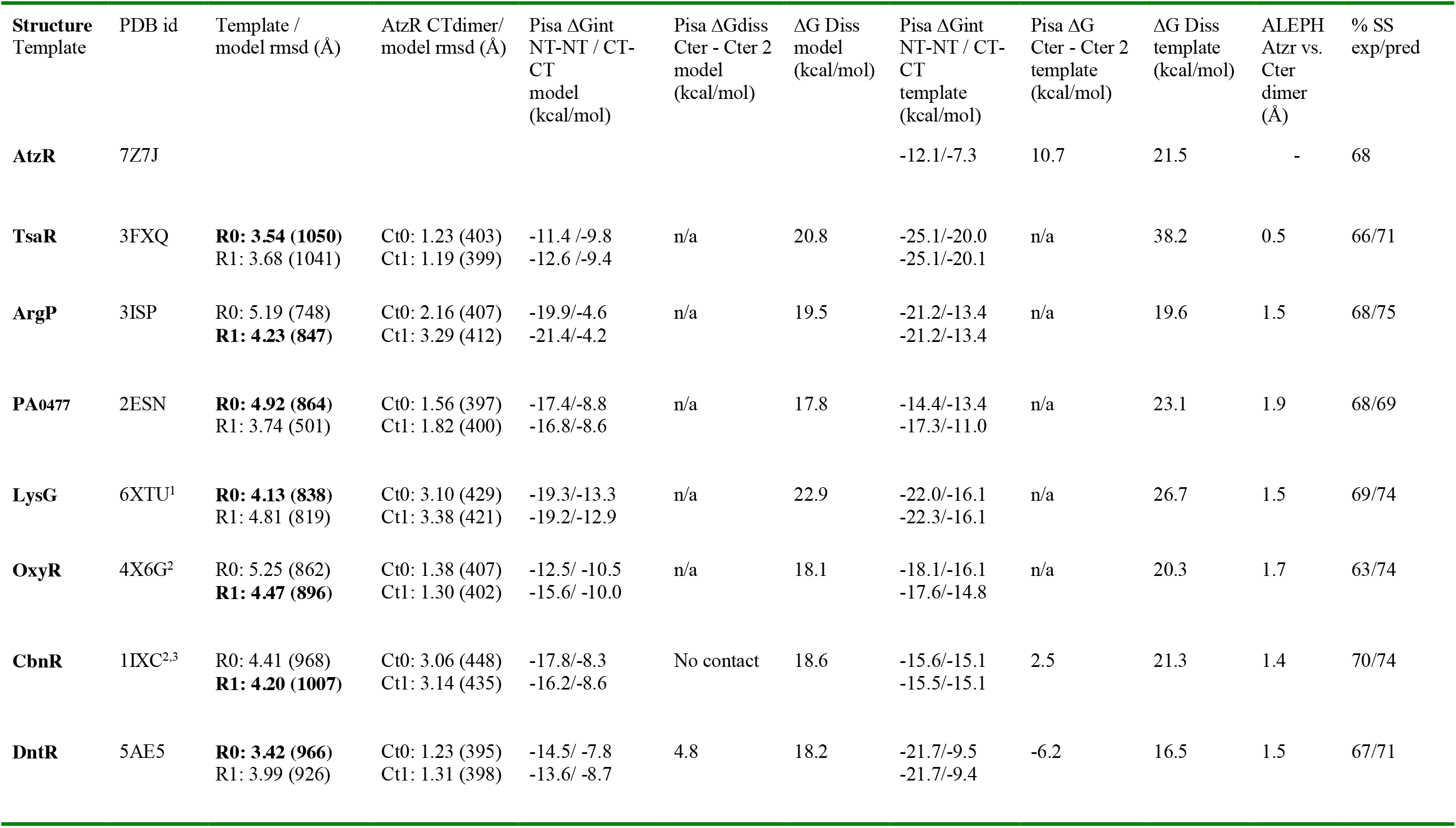
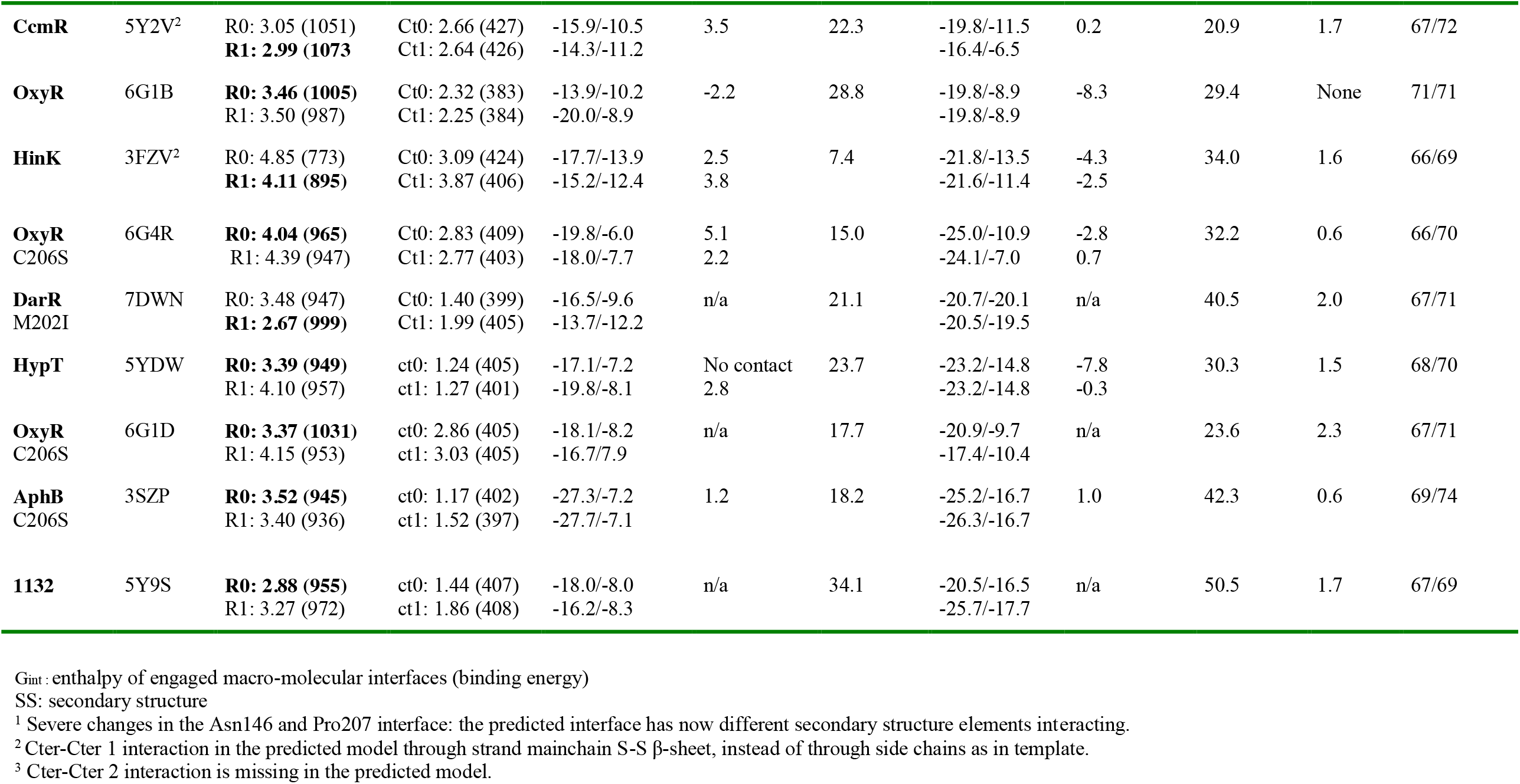
Predicted ATZR after LTTR full-length experimental structures compared with templates. Interface analysis.

The same AtzR templates used to compare known experimental structures was used to assess the dimerization interfaces in the predicted AtzR models (**Figure 3**). As can be seen from the results summarised in **Table 2**, the C-terminal dimer in all predicted structures was much closer to the one found in the experimental structure than to its templates. Superposition of the experimental dimer to predictions resulting from templates where the association between beta strands takes place through the side chains reveals larger rmsd differences. Correspondingly, the interface elements extracted with ALEPH indicate that the main chain association between strands is restored in all these predictions. Thus, the predicted effector binding dimers are closer to the one determined in the experimental structure than to their templates. Identity across the LTTR sequences is rather low, especially outside the first 80 residues. **Table 1** quotes overall values and for N-terminal domains relative to AtzR. The closest homologues in the set correspond to TsaR and CbnR, with 23 and 22% sequence identity respectively (35 and 36% for N-terminal domains). Remarkably, TsaR renders one of the closest predictions, with 3.54 Å rmsd for 1050 Cα pairs whereas the prediction for CbnR deviates more, with 4.20 Å rmsd for 1007 Cα pairs. Prediction and template are compared in **Figure 4** for TsaR. Overall agreement is high, and also for the C-terminal dimerization interface the prediction closely mimics the TsaR structure. Comparing the backbone of the effector-binding domain in the prediction versus experimental AtzR structure shows high agreement, with main deviations in the regions where interactions with the bound effector are taking place in the experimental structure. This situation contrasts with the case of CbnR (**Figure 5**), where the interface closing the ring in the template structure is missing in the prediction. This is one of the templates where the C-terminal dimers are dimerising with a side chain interaction between their beta strands. **Figure 6** shows areas with main differences in predictions from their templates. Predictions are coloured in a range representing pLDDT. In the case of LysG (PDB ID 6XTU, Della Corte *et al*., 2020), the region 200-215 forms a loop rather than the most frequently found helix. Compact monomers contact across the tetrameric ring through residues in this loop and 151-152. Still, the ring is not closed as in type 2 tetramers since the area is poorly structured, displaying comparatively high B-values. As seen in **Figure 6a**, the prediction displays in this region the alpha helix seen in the AtzR structure, and the resulting, more compact domain recedes, too far from the 151-152 loop for an interaction. The lower confidence in this predicted helix is reflected in the lower pLLDT. **Figures 6b-d** show three of the four cases where the C-terminal dimer is predicted as in the AtzR structure, with a mainchain interaction of the beta strands across subdomains, and not as in the respective templates. Despite the change from the template, pLDDT values are high in all cases, reflecting high confidence. The C-terminal dimers in the predictions turned out closer to the experimental AtzR structures than to their templates in all cases, with residual differences in the effector binding cavity as illustrated in **Figure 3c**. A prediction of the AtzR tetramer with its own experimental structure as template illustrates the sequence dependency in the template: whereas using the experimental structure renders an rmsd of 0.43Å between template and prediction, the difference amounts to 1.80 Å when all residues in the template are mutated to alanine. Main differences are found in the orientation of the mobile DNA-binding motifs and in the cyanurate binding environment displayed in **Figure 7**, where the cavity becomes narrower as the nearby chains move into the empty space.

**Figure 4.**
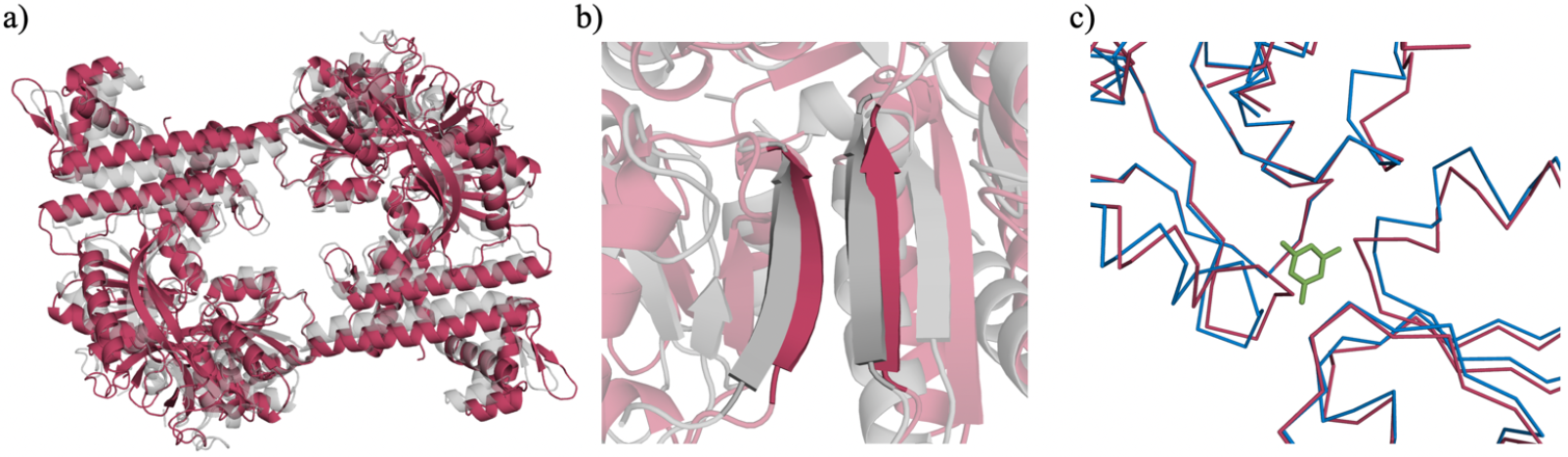
Comparison of the predicted AtzR model (red) after TsaR. **a**) Overall superposition with the TsaR template (grey). **b**) Dimerization of the C-terminal domains for prediction and template (grey). **c**) Detail of the ligand-binding area in the AtzR experimental structure (blue). In green, ligand found in experimental AtzR. The environment of the ligand showing differences is mainly S198 to G201 (left), S99 to A103 (right) and A246 to T249 (top).

**Figure 5.**
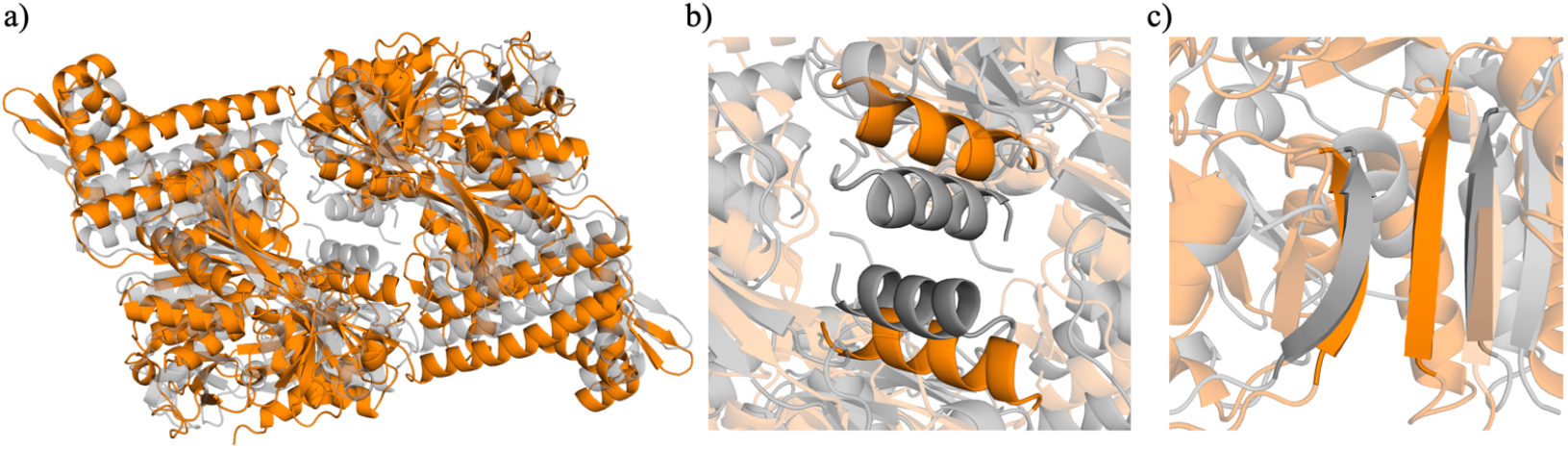
Superposition of CbnR (1IXC) used as template for the prediction (grey) and the resulting AtzR model (orange). **a**) overall models. **b**) changes in the central interface closing the tetrameric ring. **c**) changes in the beta-strands from the dimerization interface of the effector binding domains. Note that the secondary structure elements highlighted in b) and c) correspond to residues aligned in AtzR and CbnR.

**Figure 6.**
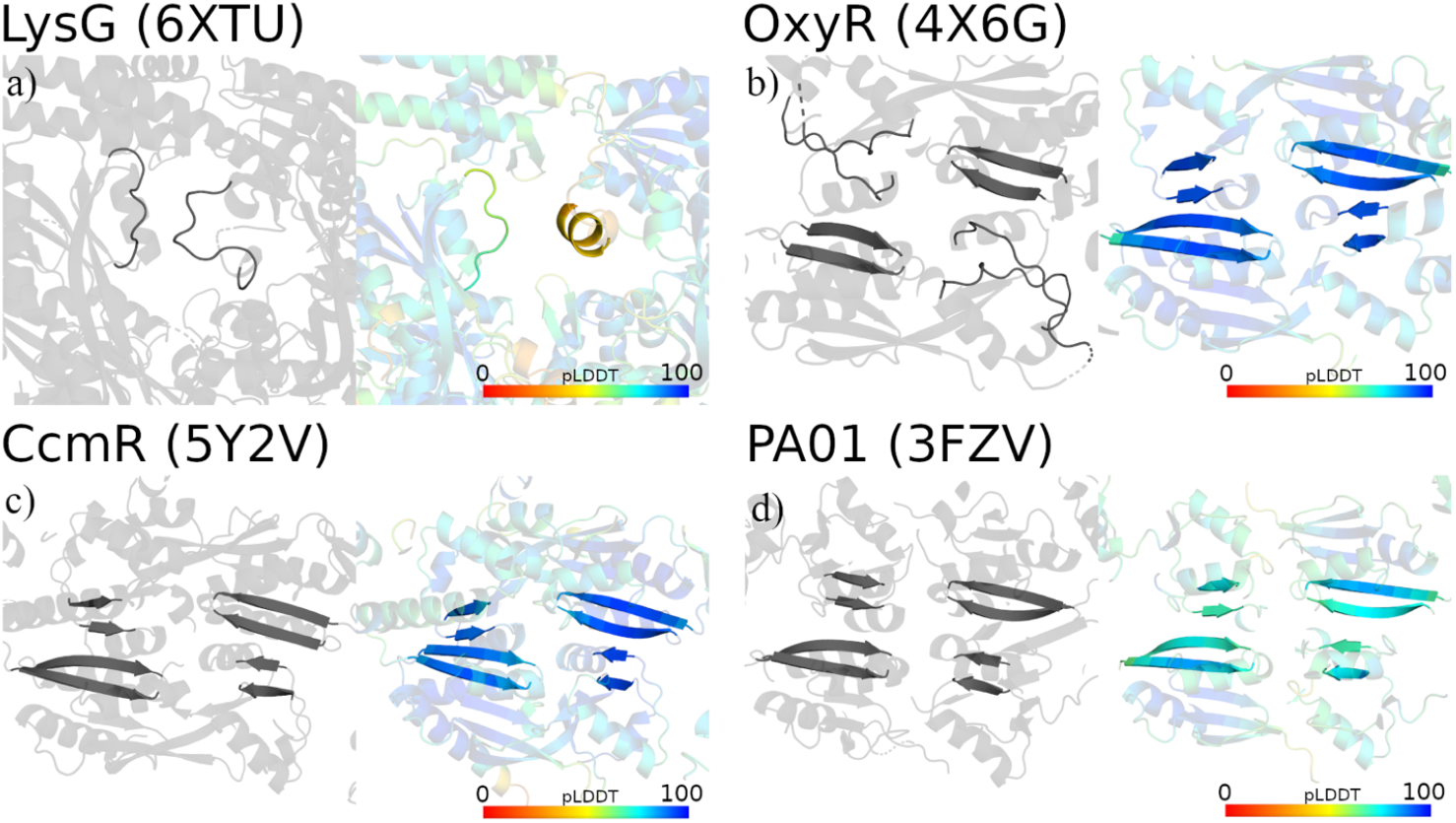
Comparison of template and prediction as cartoon in the structures and regions showing largest differences along with CbnR (1IXC displayed in Figure 5). **a**) Differences between template and AtzR-LysG in the interface involving Asn146 and Pro207 residues. **b-c-d**) Cter-Cter 1 differences between templates and AtzR-OxyR, -CcmR and -HinK predictions, respectively. Left panels: templates (in grey). Right panels: predictions (coloured according to pLDDT values).

**Figure 7.**
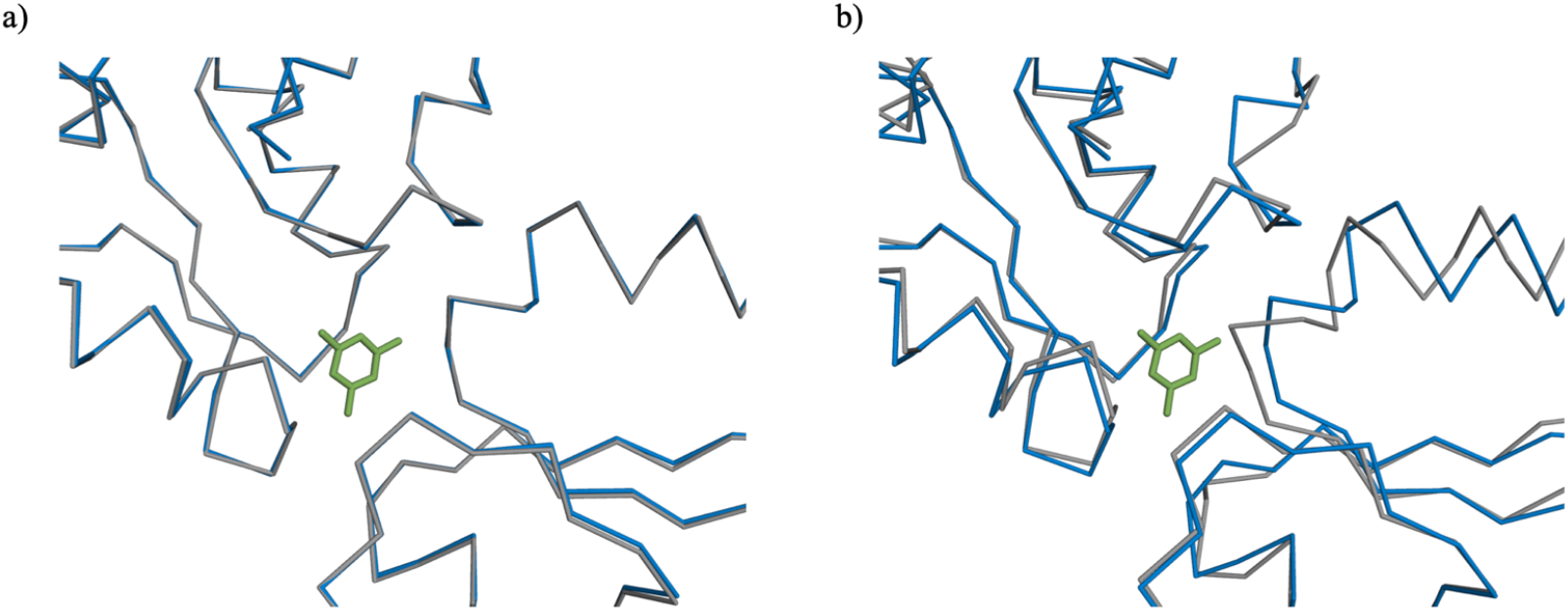
Differences in the prediction of Atzr in the environment of the effector binding site depending on the sequence of the template. Template in blue with cyanurate in green. Prediction in grey. a) the experimental AtzR with side-chains as template b) poly-alanine template.

PISA was used to calculate ΔG values characterising the monomers as well as the interfaces formed in experimental structures and in predictions. For the monomers, comparable ΔGint values ranging from -250 to -280 Kcal/mol are derived for predictions and experimental structures. Free energy gain associated to the various interfaces and dissociation energies for the tetramers are summarised in **Table 2**. It can be seen that although a higher stabilisation is always predicted for the experimental structures the values obtained are generally in a comparable range and that all predictions render tetramers expected to be stable. Nevertheless, the prediction after HinK, classified as type 3 stands out through its lowest ΔGdiss of 7.4 Kcal/mol, compared with the 34 Kcal/mol of its template. This value is also significantly lower than for all other predictions and experimental templates. Also the 4- and 5-type predictions show larger ΔGdiss gaps to their templates than the multimers in types 1 and 2. For them, estimated stabilisation of the particle does not exceed the values derived from the formation of the ring common to all structures, in spite of the additional interfaces their conformations could prompt. The experimental structure of AtzR is characterised by a remarkably extended interface closing the tetrameric ring but involving contacts not seen in any of the other LTTR.

## Discussion

The regulatory mechanism of LTTR requires dynamics that are difficult to characterise as the information comes from different species and sample stabilisation is seen to dominate over the effect of interactions with effectors or DNA. In the present work we have used AF to relate structures from different species by predicting versions of the AtzR structure mimicking the conformations seen in the diverse experimental structures and comparing them to our crystallographic determination of the full-length AtzR in complex to its effector, cyanuric acid.

Interestingly, predictions for the C-terminal domain dimer resembles more the experimental structure than the templates and structures like the one predicted after TsaR show their main local differences in the ligand-binding environment. These differences suggest an intrinsic preference for the association mode seen in the dimer, excluding a rotation between C-terminal domains as part of the dynamic. These results might be prompted by the tendency observed towards more ordered, canonical structure as shown by the higher percentage of regular secondary structure in the predictions than in the experimental templates. On the other hand, preliminary tests show that the CbnR prediction after the TsaR template reproduces its own experimental side chain association rather than adopting the sheet extension seen in TsaR (data not shown).

It is also noteworthy that whereas all predictions show favourable energy for the tetrameric particles mimicked and the less constrained type 1 conformations and some type 2 are reasonably plausible, in two cases (CbnR- and HypT-like) the second C-terminal contact is absent in the prediction and the conformation has drifted enough to avoid the additional interfaces in the template. In some cases, the energy differences are such that trial conformations could be ruled out. Exemplarily, the much lower ΔGdiss value shown for the very compact HinK conformation would suggest that the required interfaces are unlikely to be formed as targeted in the prediction and that the conformation forces distortion on the main interfaces. All of the type 4 and 5 tetrameric structures appear to fall short from their templates. As mentioned, it can be observed that the fraction of regular secondary structure tends to be higher in all the predictions than in the experimental LTTR structures. It is remarkable that some of the LTTR conformations show poorly structured regions in the hinge and beginning of the linker helix, maybe a structural price to pay to achieve the particular tetrameric conformation and an energy trade off in the mechanism. AtzR itself shows some poor electron density in its N-terminal domains as does DntR, both feature rather extended contacts in the additional C-terminal interface closing the ring. This may be a reason why closest predictions correspond to templates of more ideal secondary structure, for instance TsaR.

## Concluding remarks

We have modelled dynamics with AF to analyse the new AtzR crystallographic structure in the context of known structural data. Being able to use AF in a flexible homology-modelling mode to produce heterotetrameric models may help in understanding the transition between conformations that are triggered by effector binding in case correspondence between an open and closed state underlies all systems and here we have probed the method with a new structure outside the AF training set. Dissociation energy estimated for the predicted multimers discourage the hypothesis that each LTTR would adopt all the conformations seen. Currently, potential uses of predictions raise more questions than they can settle.

The method implemented in ARCIMBOLDO_AIR to examine the structural experimental evidence considering dynamics in the context of predictions can be used for other systems. Sequence identity between templates and predictions opens a way to selectively constrain structure conferring targeted degrees of freedom and setting boundary conditions. This is indicated by the effect of poly-alanine templates versus final residue composition and will be further explored. In any case, developing tools to integrate predictions and biochemical and structural data should allow to integrate knowledge broadly across experimental cases.

## Acknowledgements

This work was supported by grants PGC2018-101370-B-100 and BIO2007-63754 (MICINN/AEI/FEDER/UE) and STFC (CCP4-ARCIMBOLDO_LOW). AM is grateful to MICINN for her BES-2017-080368 scholarship associated with the Structural Biology Maria de Maeztu Unit of Excellence (MDM2014-0435-01). Support from STFC/CCP4 is gratefully acknowledged. We thank Rafael Borges for helpful discussion.

---

**Table 1Extended.** Summary of crystallographic structures of LTTR effector binding domain dimers. Analysis of interfaces and comparison with the corresponding interface in the AtzR structure if present.

| Structure Template | PDB id | Multimer type | Cter-Cter 1<br>113-235/130-225 | Cter-Cter 2 | ALEPH AtzR<br>vs. Cter dimer |
| --- | --- | --- | --- | --- | --- |
| **AmpR**<br>*C. freundii* | 4WKM | Functional<br>Cterm dimer | H-H, S-S beta-sheet, twisted | – | None |
| **BenM**<br>*A. baylyi* | 2F6G (2F6P, 2F78, 2F7A, 2F8D, 2F97, 2H99, 2H9B) | Functional<br>Cterm dimer | No interface, missing sheet | 2F8D (benzoate); 2F97: H200-215 (~CbnR, DntR) | None |
| **CatM**<br>*A. baylyi* | 2F7B (2F7C, 3GLB, 2H98) | Functional<br>Cterm dimer | No interface, missing sheet | H200-215 (~CbnR, DntR) | None |
| **CcpC**<br>*B. amyloliquefaciens* | 7DMW | Functional<br>Cterm dimer | H-H, S-S beta-sheet |  | 1.6 Å |
| **CmpR**<br>*S. elongatus* | 5Z49 | Functional<br>Cterm dimer | H-H, S-S beta-sheet + H-H, S-S sidech. | 150, 200 (~CbnR, DntR) | 2.6 Å |
| **CrgA**<br>*N. meningitidis* | 3HHF | Functional<br>Cterm dimer | H-H, S-S beta-sheet | – | 1.9 Å |
| **CysB**<br>*S. typhimurium* | 4LON (4GWO, 4LP2, 4LQ2, 4LQ5, 4M4G) | Functional<br>Cterm dimer | H-H, S-S beta-sheet, twisted | – | None |
| **DntR**<br>*Burkholderia* | 2Y84 (2Y7K, 2Y7P, 2Y7R, 2Y7W, 5AE4, 2UYF, 2UYE) | Functional<br>Cterm dimer | H-H, S-S beta-sheet | 2UYE, 2UYF: H200-215 (~CbnR, DntR) | 1.8 Å |
| **HypT**<br>*S. typhimurium* | 5YDV (5YDO, 5YER, 5YEZ) | Functional<br>Cterm dimer | Tetramer like<br>H-H, S-S beta-sheet | – | 1.5 Å |
| **LeuO**<br>*E. coli* | 6GZ0 (6GZ1, 6GZ2) | Functional<br>Cterm dimer | H-H, S-S beta-sheet | – | 2.7 Å |
| **OccR**<br>*A. tumefaciens* | 5VVI (5VVH) | Functional<br>Cterm dimer | H-H, S-S beta-sheet | – | 2.0 Å |
| **OxyR**<br>*P. aeruginosa* | 4Y0M (3JV9, 4XWS) | Functional<br>Cterm dimer | No interface, missing sheet | – | None |
| **OxyR2**<br>*V. vulnificus* | 5X0V (5B7H, 5B70, 5B7D, 5X0Q) | Functional<br>Cterm dimer | No interface, missing sheet | – | None |
| **PqsR**<br>*P. aeruginosa* | 4JVC (4JVD, 4JVI, 6YIZ, 6Q7U, 6Q7V, 6Q7W, 7NBW) | Half domain interaction | Half interface | – | None |
| **QuiR**<br>*L. monocytogenes* | 5TED | Functional<br>Cterm dimer | H-H, S-S beta-sheet | – | 2.9 Å |
| **RovM**<br>*Y. pseudotuberculosis* | 3ONM | Functional<br>Cterm dimer | H-H, S-S side-chains | – | None |

\* Geometric comparison for isolated C-terminal dimers: an AtzR-like interface is found in no contacts but some cases display an interface similar to CbnR or DntR.

## METHODS

### Test cases of LTTR

Test cases were selected from all deposited crystallographic structures of full-length LTTR in non-redundant, isostructural conformations, even those where the DNA-binding region is not exposed on the surface of the oligomer, although their participation in the transcription regulation mechanism is difficult to envisage.

AtzR (PDB ID 7Z7J), TsaR (PDB ID 3FXQ, Monferrer *et al*., 2010), ArgP (PDB ID 3ISP, Zhou *et al*., 2010), PA0477 (PDB ID 2ESN), LysG (PDB ID 6XTU, Della Corte *et al*., 2020), OxyR (PDB ID 4×6G, Jo *et al*., 2015), CbnR (PDB ID 1IXC, Muraoka *et al*., 2003), DntR (PDB ID 5AE5, Lerche *et al*., 2016), CcmR (PDB ID 5Y2V, Jiang *et al*., 2017), OxyR (PDB ID 6G1B, Pedre *et al*., 2018), HinK (PDB ID 3FZV, Wang *et al*., 2021), OxyR (PDB ID 6G4R, Pedre *et al*., 2018), DarR (PDB ID 7DWN, Wang *et al*., 2021), HypT (PDB ID 5YDW, Jo *et al*., 2019), OxyR (6G1D, Pedre *et al*., 2018), AphB (3SZP, Taylor *et al*., 2012), 1132 (5Y9S, Jang *et al*., 2018) and CrgA (3HHG, Sainsbury *et al*., 2009)

Details on the data and tests are summarized in **Table 1** and **Table 2**.

Models produced have been deposited under https://gitlab.com/arcimboldo-world/af-lysr-regulators

### Structure determination of AtzR at 1.85 Å

For heterologous production of AtzR in *Escherichia Coli* M10, 500 mL TB media (4 mLL^−1^ glycerol, 12 gL^−1^ peptone, 24 gL^−1^ yeast extract, 0.17M KH_2_PO_4_, 0.74M K_2_HPO_4_) supplemented with 50 mgL^−1^ kanamycin was inoculated using 10% (v/v) of the respective overnight culture, and protein expression was directly induced by adding 0.2M cyanuric acid. After 2-h incubation (20°C, 200 r.p.m.), cells were harvested by centrifugation (4400g, 20 min at 4°C) and cell pellets were stored at −20°C until further processing. The pellet obtained from 2 L cell culture was resuspended in 40 ml lysis buffer (20 m*M* Tris–HCl pH 7.0, 0.3 *M* NaCl) containing either 200 µ*M* PMSF or a tablet of complete EDTA-free protease-inhibitor cocktail (Roche) and 10 ng ml−1 DNAse I. Cells were lysed using a Cell Disruptor (Constant Systems Ltd) applying a pressure of 1.35 MPa and centrifuged for 30 min at 40 000*g*. Purification of resulting protein was performed by immobilized metal affinity chromatography using a 5 mL HisTrap HP column (GE Healthcare, Frei-burg, Germany) containing HisTrap chelating stationary phase (Amersham Biosciences) with 5 mL bed volume. Unbound proteins were removed by first eluting using a washing buffer with imidazole (20mM Tris pH7.5, 0.5M NaCl, 5% Glycerol, 300mM Imidazole and 2.25mM TCEP). The protein was eluted with the same buffer but with an increased concentration (500 mM) of imidazole. The purified protein was concentrated up to 5 mg/mL using Centricon YM-10 and Microcon YM-10 concentrators (Amicon, Millipore). Crystals used in this study were obtained under oxygen exclusion in hanging drops from 0.5 M Hepes pH 7.2, although they can be produced at pH 7 to 8 and alternatively with 0.5 M EPPS as buffer.

Data collection, 900 frames of 0.1° oscillation range were collected from a single crystal at the Proxima (Soleil) beamline at a wavelength of 0.98011 Å. The data were integrated using the program XDS (Kabsch, 2010) and further processed through the autoPROC (Vonrhein *et al*., 2011) expert system, which employs AIMLESS (Evans & Murshudov, 2013) from CCP4 (Winn *et al*., 2011) for scaling, and StarAniso (Tickle *et al*., 2018-2021) for elliptical truncation in case of anisotropy. This resulted in data truncated to 2.22Å along a* and b*, and to 1.80Å along c*. The unique data had a completeness of 94.3% in the corresponding ellipsoid in reciprocal space, and a mean multiplicity of 12.6.

Data characteristics are summarised below:

|  |  |
| --- | --- |
| PDB ID | 7Z7J |
| Resolution (Å) | 77.7-1.80 (1.96-1.80) |
| R<sub>meas</sub> | 0.044 (1.09) |
| CC<sub>1/2</sub> | 1.00 (0.83) |
| <I/σ<sub>I</sub>> | 27.3 (2.3) |
| Number of observations | 525047 (19680) |
| Number of unique reflections | 41572 (2079) |
| Multiplicity | 12.6 (9.5) |
| Percent completeness (spherical) | 65.3 (14.7) |
| Percent completeness (elliptical) | 94.3 (84.9) |
| R factor (%) overall (1.85-1.8) | 24.1 (35.0) |
| Free R factor (%) overall (1.85-1.8) | 28.5 (45.8) |

The structure was solved with ARCIMBOLDO (Millán *et al*., 2015) using AlphaFold2 models to derive fragments, place them with Phaser (McCoy *et al*., 2007) and expand partial solutions with SHELXE (Usón & Sheldrick, 2018). The structure was refined with Refmac (Murshudov *et al*., 2011), TLS groups were defined separating the DNA-binding motif, linker helix and effector binding domain. COOT (Emsley & Cowtan, 2004) was used for real space refinement and manual building.

### Prediction with AF

Predictions run for two to three hours on a workstation of the following characteristics: AMD Ryzen Thread Ripper 3975WX, Nvidia GeForce RTX 3090 24 GB. AF was run on a virtual machine using 48 out of the host’s 64 cores, and 192 out of its 256 GB RAM, the OS of the virtual machine was Ubuntu 20.04.4 LTS. AF2 code downloaded from DeepMind’s GitHub repository was modified in order to skip all the default steps from the Data module and read a set of custom features for the inference and store them in a pickle file. Four copies of the sequence were concatenated inserting linker stretches of 50 glycines. The length of this stretch was originally estimated from distances in the templates but its particular value is not relevant. All the predicted models were calculated in the monomeric mode of AF2 using tetrameric templates from the experimental structures. In those cases where tetramers were not present in the ASU, symmetry operations were applied to build the functional tetrameric assembly. Template information was introduced in the features.pkl file minding the gaps in the numbering to match the poly-glycine stretches inserted in the sequence. Best predicted models were assessed by computing the superposition to its corresponding template with Superpose from CCP4 suite (Winn *et al*., 2011).

### Assessment and comparison

Multimer interfaces were compared and classified across experimental and predicted tetramers and dimers with ALEPH using a template composed by the secondary structure elements involved in contacts building the C-terminal dimerisation interface. The AtzR regions Glu105-Ala113, Ala122-Lys126, Gln219-Ser222 and Met227-Glu234 were extracted from both monomers in the crystallographic structure. These fragments were used as a template to perform a geometrical search and superposition based on *characteristic vectors* with ALEPH against all the experimental and predicted models. Chain assignment filters were applied on the extracted fragments to ensure that the interface was located and the lowest rmsd between template and extracted local fold was selected. Results were manually curated to exclude artefacts upon failure to extract the interface.

### Energy assessment in PISA

All predictions were treated as isolated tetrameric particles in solution by increasing the unit cell in *P*1 symmetry for subsequent energy calculations with PISA (Krissinel, 2011). For energy calculations between interfaces in monomers, predictions were split in different chains and poly-glycine linkers were removed.

